# Linking parasitism to network centrality and the impact of sampling bias in its interpretation

**DOI:** 10.1101/2021.06.07.447302

**Authors:** Zhihong Xu, Andrew J. J. MacIntosh, Alba Castellano-Navarro, Emilio Macanás-Martínez, Takafumi Suzumura, Julie Duboscq

## Abstract

Group living is beneficial for individuals, but also comes with costs. One such cost is the increased possibility of pathogen transmission because increased numbers or frequencies of social contacts are often associated with increased parasite abundance or diversity. The social structure of a group or population is paramount to patterns of infection and transmission. Yet, for various reasons, studies investigating the links between sociality and parasitism in animals, especially in primates, have only accounted for parts of the group (e.g., only adults), which is likely to impact the interpretation of results. Here, we investigated the relationship between social network centrality and an estimate of gastrointestinal helminth infection intensity in a whole group of Japanese macaques (*Macaca fuscata*). We then tested the impact of omitting parts of the group on this relationship. We aimed to test: (1) whether social network centrality – in terms of the number of partners (degree), frequency of interactions (strength), and level of social integration (eigenvector) – was linked to parasite infection intensity (estimated by eggs per gram of faeces, EPG); and, (2) to what extent excluding portions of individuals within the group might influence the observed relationship. We conducted social network analysis on data collected from one group of Japanese macaques over three months on Koshima Island, Japan. We then ran a series of knock-out simulations. General linear mixed models showed that, at the whole-group level, network centrality was positively associated with geohelminth infection intensity. However, in partial networks with only adult females, only juveniles, or random subsets of the group, the strength of this relationship - albeit still generally positive - lost statistical significance. Furthermore, knock-out simulations where individuals were removed but network metrics were retained from the original whole-group network showed that these changes are partly a power issue and partly an effect of sampling the incomplete network. Our study indicates that sampling bias can thus hamper our ability to detect real network effects involving social interaction and parasitism. In addition to supporting earlier results linking geohelminth infection to Japanese macaque social networks, this work introduces important methodological considerations for research into the dynamics of social transmission, with implications for infectious disease epidemiology, population management, and health interventions.

## Introduction

Living in a group provides many benefits to individuals, but also substantial costs (Loehle, 1995). One such cost is increased exposure to parasites and a higher probability of parasite transmission from one individual to another (Freeland, 1976; Loehle, 1995; Arneberg et al., 1998; Nunn et al., 2003). This pattern may arise from higher contact rates between individuals and generally higher local densities of hosts. From the perspective of vectors of parasites, a group of animals is an aggregation of hosts, potentially more attractive than a single host (Arneberg, 2002; Nunn et al., 2003). Even for parasites that are not transmitted through direct contact between hosts, sharing of resources and space, such as food patches, sleeping sites, or home ranges, increases the likelihood that hosts encounter infectious stages of parasites in the environment, which increases the probability of transmission between hosts (Altizer et al., 2003; Nunn et al., 2015; Müller Klein, M. Heistermann, et al., 2019).

Although living in a group may generally mean greater parasite exposure, within a group, individuals exhibit different rates of interacting: social interactions are influenced by kinship ties, dominance ranks, or age, size, and sex assortativity (Kurvers et al., 2014). This creates heterogeneity in contact patterns, leading to differences between potential contacts and actual contacts, thereby influencing each individual’s exposure to parasites (Altizer et al., 2003). Thus, the positive relationship that is expected to occur between increased group size and increased parasitism is not always observed (Moller et al., 1993). Accounting for social structure and the role and position of individuals within it therefore helps make sense of how parasites are transmitted within animal groups. Studying the relationship between social interactions and parasite infection is essential for understanding how animal social behaviours might reflect adaptations to infection risk, or conversely how parasites might invade or persist in host populations.

In this endeavour, Social Network Analysis (SNA) is an efficient statistical tool that can be used to analyse groups of individuals as a whole and generate predictions about the spread of infectious diseases (Borgatti, 2005). Social networks reflect the patterns of interactions, associations, or spatial proximity between individuals. SNA can provide information about the whole structure of the network, but also about the structural importance of certain nodes (e.g. individuals) and their ties (e.g. interactions) to identify influential individuals within a network (Borgatti et al., 2009). So-called central individuals have qualitatively or quantitatively greater direct or indirect connections than less central or peripheral individuals. Central individuals are thus expected to be at once key dispersers of parasites but also at the greatest risk of acquiring parasites from others (Romano et al., 2016). For example, gidgee skinks (Egernia stokesii) with high refuge-sharing rates are more likely to be infected by ticks and host a more diverse blood parasite community than less-connected skinks in the same refuge-sharing network (Godfrey et al., 2009). Similarly, female Japanese macaques (Macaca fuscata yakui) that are better connected in their grooming networks exhibit more diverse intestinal helminth communities and greater intensity of infection with certain parasite species (MacIntosh et al., 2012). In cases such as the latter, carefully designed experiments are required to test the mechanisms linking networks to infection. But in general, such links have been demonstrated in a broad array of contexts, regardless of the type of network or parasite species under study, or even the measures of parasitism used (Altizer et al., 2003; Ezenwa et al., 2016; Rushmore et al., 2017; Briard and Ezenwa, 2021; Lucatelli et al., 2021).

Factors like sex, social rank, and age can also influence host exposure and susceptibility to parasites. These traits might have an impact on the frequency, quality, and quantity of interactions with social partners and with the environment. For instance, in species where young males disperse from their natal group, males will interact with more a numerous and diverse set of individuals compared with females (Teichroeb et al., 2011). In addition, these traits might be linked to variations in physiological and immunological responses. As a case in point, meta-analyses in vertebrates suggest that greater parasitism is biased toward high-ranking individuals (Habig et al., 2018). This may be linked to stressful social situations - such as the struggle to achieve high dominance status - that trigger the production of glucocorticoids, which is essential in the activation of the immune response (Cain and Cidlowski, 2017). At the same time, however, dominance status can also correlate with contact networks, making it hard to disentangle the mechanisms of infection using observational data (MacIntosh et al., 2012; Wooddell et al., 2020).

Concerning age, juveniles may have a higher chance of contacting infectious agents because of their specific activity patterns, which means higher exposure to infection (e.g., greater and more diverse exploration of substrates, heightened reliance on social interactions, etc.) (Sarabian, et al., 2018). Similarly, juveniles are known to have a less efficient or weaker immune system than adults, which increases their susceptibility to infection. (Fallon et al., 2003; East et al., 2008; Ebersole et al., 2008). Often, though, studies investigating links between sociality and parasitism only include adults, perhaps because conducting research on juveniles is more difficult due to more challenging identification and observation conditions. Many studies, including those by some of us (MacIntosh et al., 2012; Duboscq et al., 2016; Romano et al., 2016; Webber et al., 2016; Hamilton et al., 2020; Roberts and Roberts, 2020; Sandel et al., 2021), have omitted juveniles, and this may preclude a fuller understanding of infection dynamics. As a case in point, although unrelated to infection, a previous study using knockout simulations indicated that juvenile or adolescent baboons can act as bridges between clusters of otherwise weakly connected individuals, thus changing the nature of the networks being observed (Fedurek and Lehmann, 2017).

More generally, problems of missing data in social networks are well known. Missing nodes (Kossinets, 2006; Smith and Moody, 2013; Davis et al., 2018; Gilbertson et al., 2021), especially missing central nodes (Smith et al., 2017), produce measurement bias in social network centrality because network data are relational and hence one data point (node, edge, etc) is not independent of another. Although many network measures can be estimated with incomplete information and can be robust to missing individuals (Borgatti et al., 2006), it is important to assess the impact on precision, accuracy, and bias created by subsampling networks or by carrying out analyses with partial networks (Silk et al., 2015; Silk, 2018). This impact can be accessed via comparing, correlating, or regressing original whole-group network measures with newly calculated partial network measures. At present, we are starting to better understand the effects of using partial or subsampled networks on network measures (Silk et al., 2015; Smith et al., 2017), but we have scarce knowledge about the influence that such measurement bias has on the conclusions we might draw concerning the relationship between network characteristics and other social or ecological processes (but see Silk et al., 2015 or Herrera et al., 2021). Assessing measurement bias can thus inform us about how best to carry out our observations or even how we might correct for sampling bias when data have already been collected (Silk et al., 2015; Smith et al., 2017; Hoppitt and Farine, 2018). It is thus meaningful to assess how including (or excluding) juveniles or any other often-missing subgroup(s), or even individuals at random, might affect our interpretation of the relationship between infection and social network centrality, which is the target of this study.

Japanese macaques and their gastrointestinal parasites comprise a well-suited system for studying pathogen transmission in wildlife (MacIntosh, 2014). Japanese macaque populations are generally stable, are distributed throughout Japan, and have been subject to a long history of research, which provides a firm foundation regarding their ecology and social structure. Japanese macaques are mainly terrestrial, enabling direct observation of social behaviour and sampling for parasitological analyses. Japanese macaques are also infected by numerous gastrointestinal nematode parasites, including various geohelminths, which are amenable to study through faecal sampling (Gotoh, 2000). Helminths are commonly found in mammals (Stephens et al., 2017) and can impact host health and fitness by imposing nutritional constraints and other physiological conditions that can reduce host fitness, and in some cases may directly cause mortality (Hillegass et al., 2010).

There are three known geohelminths in Japanese macaques: *Oesophagostomum aculeatum* (Strongylida), *Strongyloides fuelleborni* (Rhabditida), and *Trichuris trichiura* (Trichocephalida) (Gotoh, 2000). These parasites are transmitted through contact with contaminated substrates, though while *O. aculeatum* and *T. trichiura* use the faecal-oral pathway, *S. fuelleborni* infects hosts percutaneously. Voided with faeces, the eggs of *O*.*aculeatem* and *S. fuelleborni* hatch and develop into infectious third-stage larvae in the environment. *S. fuelleborni* exhibits a heterogonic life cycle whereby some larvae continue to develop in the external environment and persist as free-living adults, mating and reproducing opportunistically. Others (females only) become parasitic in a host. In contrast, the eggs of *T. trichiura* become embryonated but do not hatch until they have been ingested by a suitable host. Each can thus be transmitted between hosts that share environmental resources, even if separated in time (Modrý et al., 2018).

The present study examined gastrointestinal nematode parasites infecting Japanese macaques on Koshima island, Japan, to test two hypotheses: (1) to what extent social network centrality, as measured by the number of partners (degree), frequency of interactions (strength), and social integration (eigenvector) observed across individuals, was linked to an estimate of intestinal parasite infection intensity (eggs per gram of faeces, EPG); and, (2) to what extent analyses based on partial networks produced different results than analysis based on a whole-group network. First, we predicted a positive association between an individual’s social network centrality and its gastrointestinal nematode parasite infection intensity (estimated via faecal egg counts), as was found previously in Japanese macaques (MacIntosh et al., 2012) and other primates and environmentally-transmitted parasites (Grear et al., 2013; Friant et al., 2016; Wren et al., 2016; Müller Klein et al., 2019). We chose multiple network metrics because, in theory, each may be associated with a different aspect of transmission or may be more or less instrumental to the process (MacIntosh et al., 2012; Duboscq et al., 2016; Tiddi et al., 2019). Second, we predicted that accounting for only partial networks would lead to divergent results concerning the network-infection link. Partial networks examined included a juvenile-only network, because of their high rates of infection (MacIntosh et al., 2010), an adult female-only network, because of their core role in macaque society and their prominence in previous studies of this nature, and a series of random subsets of the network. We do not make more precise predictions here as we cannot a priori predict with confidence the effects of missing individuals on network measures.

## Methods

### Ethical Statement

The research presented here complied with the Guidelines for the Care and Use of Nonhuman Primates established by the Primate Research Institute of Kyoto University (KUPRI), to the legal requirements of Japan, and to the American Society of Primatologists (ASP) Principles for the Ethical Treatment of Non-Human Primates. Permissions were acquired from the Field Research Committee of KUPRI, the Cooperative Research Program of the Wildlife Research Center of Kyoto University, and the City of Kushima’s Agency for Cultural Affairs.

### Study site and subjects

This research was conducted on Koshima island, a 0.35km^2^ island located in the Sea of Hyūga in Miyazaki Prefecture, Japan (31°27′N, 131°22′E). The island is inhabited by two groups of macaques and some solitary males. The home ranges of the two groups only overlap at their edges, and macaques from different groups were never observed in proximity during the observation period. The study group, the “main” group, inhabits the western part of the island, including a sandy beach, a range of forest, and a rim of rocky beach. Provisioning and behavioural observations of Koshima macaques started in 1952, and demographic, ecological, behavioural, and life-history data are available since then (Watanabe, 2008). Previous studies on this island identified four nematode parasites infecting Koshima macaques (Horii et al., 1982; Gotoh, 2000). The study group is currently provisioned with approximately 3 kg of wheat twice weekly. At the time of the study, the main group had 47 stable group members and 5 solitary roaming males that ranged in proximity to the group on an irregular basis. The group included 20 adult females (≥5 years old), 8 adult males (≥5 yo, including the 5 solitary males), 13 juvenile females (between 1 and 4 yo), and 11 juvenile males (between 1 and 4 yo). Because the population has been monitored for decades, we have the exact age of each individual in the population.

### Behavioural Data Collection

The observation team - including ZX, AC-N, EM-M - observed the macaques for 88 days over 3 months from March to June 2017. All individuals could be identified reliably based on a combination of facial and other physical characteristics (such as facial tattoos, scars, limps, coat colour, etc.), and allowed researchers to approach within 5m, facilitating observation and faecal sample collection. We carried out 20-minute focal animal observations, recording behavioural activities of the focal individual (including resting, feeding, locomotion, playing, giving and receiving grooming, and self-grooming) at 30-second intervals. At the same scan point, we also recorded the identity of each individual within 1m of, or in body contact with, the focal animal. Agonistic interactions of all individuals in the group were recorded ad libitum. To assess the dominance hierarchy, we used all agonistic interactions recorded by the observation team between November 2016 and June 2017. We observed every individual in the group, including two infants that were born in 2016, every day in a pseudorandomized order, updated daily. We avoided repeating observations of the same individual on the same day, and we balanced observations between individuals as much as possible across days and times of day (e.g., early and late morning, early and late afternoon).

Data were collected either on paper or using a tablet (Panasonic Tough-pad, with the application PRIM8 (McDonald and Johnson, 2014)). A multi-observer reliability test based on Fleiss’ Kappa index (Fleiss and Cohen, 1973) indicated a good match between observers for proximity records, including the identities of all subjects involved (Kappa = 0.815). In total, we conducted 547 focal observations recording 22,066 focal scans, or 183.3 hours of focal observations, with a mean ± SD of 10.7 ± 6.6 focal follows per individual (or 11.2 ± 0.8 per individual without 4 rarely seen solitary roaming males). Table SI-1 in the Supplementary Information summarizes the attributes of and data from the study subjects.

### Social data analysis

#### Dominance

To compute the dominance rank of all individuals, we used the Elo-rating method (Neumann and Kulik, 2014), which reflects an individual’s success in agonistic interactions relative to all other members (Albers and de Vries, 2001; Neumann et al., 2011). Elo-rating is based on a sequence of agonistic interactions ordered in time. Each individual is seeded with an arbitrary rating of 1000. This rating is then updated after each agonistic interaction with a clear outcome. The rating increases/decreases based on the outcome of the interaction (won or lost), the previous ratings of both opponents, and a determined factor, k (here k = 100 (Neumann et al., 2011)). In our analyses, we aggregated all data and calculated Elo-ratings for all individuals at the end of the observation period.

#### Social network centrality

We constructed an undirected proximity network based on recorded 1m (close) proximity. The parasites of interest in this study are generally thought to be transmitted primarily through contact with contaminated substrates. In the context of group dynamics, resource sharing and shared space use rather than direct social contact might be implicated in the transmission of such parasites. We thus assumed that a close-proximity network was representative of the probability of (in)direct transmission-relevant space or resource sharing among individuals in this highly social species. Our previous work with Japanese macaques, as well as various other studies of other species, support this assumption (MacIntosh et al., 2012; VanderWaal et al., 2013; Friant et al., 2016; Müller Klein, et al., 2019).

We constructed a network based on aggregate data over the whole study period (3 months). It’s difficult to assess how representative of transmission processes this aggregate network might be. Data from livestock suggest prepatent periods – i.e., the period between acquisition of a parasite and the point at which it (if female) begins shedding eggs in host faeces – of between 2 and 6 weeks for the parasite genera studied here (Anderson, 2000). Thus, networks based on shorter time periods may better match the time lag between infection and detection. However, such networks, given sampling limitations, would not be robust representations of connections between our study subjects. We recognise the limitations of using aggregate behavioural data over arbitrary time scales to infer transmission processes, and so we instead emphasize that our network data should be seen as reflecting generalized patterns of interaction that can be related to observed infection phenotypes of group members – themselves as an aggregate of sampling over the same study period. This is an approach that appears regularly in the literature (Rimbach et al., 2015; Balasubramaniam et al., 2016; Tiddi et al., 2019), as transmission events themselves are far harder and in some cases currently impossible to study effectively. Moreover, our focus is on the phenomenological link between networks and infection and the impact of sampling variability on such links, rather than on examining the mechanics of why the link exists in the first place, so we believe our analyses are appropriate for this endeavour.

We built a symmetrical matrix of undirected social proximities, in which dyadic proximity data were scaled by the number of scans collected from each individual of the corresponding dyad. The proximity matrix was then analysed with the package “igraph” (Csardi and Nepusz, 2006) in R Statistical Software v.4.0.5 (R. Development Core Team, 2021). For each individual, we computed: (1) degree (how many partners an individual had); (2) strength (how many times an individual was in proximity with others); and (3) eigenvector centrality (the sum of the centralities of an individual’s neighbours; high centrality can be achieved by having either a large degree or being connected to associates with a high degree (or both)) (Farine and Whitehead, 2015).

### Parasite Collection and Analysis

We opportunistically collected between 1 and 4 fresh faecal samples from each individual (3.5±1.6) during observations whenever a clearly identified individual was seen defecating. We put the entire faeces into a sealable plastic bag and then into a cooler bag with ice until we brought all samples to the lab for processing. Within 12h of collection, we weighed and homogenized about 1-2 grams of faeces into 3.5ml of a sodium acetate-acetic acid-formaldehyde (SAF) solution for the preservation of parasite eggs and larvae. After the fieldwork period, we sent all samples to Kyoto University’s Primate Research Institute, where we conducted further analysis.

We processed all parasitological samples following previous work with Japanese macaque parasites using a modified formalin-ethyl acetate sedimentation protocol to concentrate helminth eggs from faeces (MacIntosh et al., 2010). We added 5 drops of Triton X-100 to the faecal mixture before filtration to better isolate eggs from other components of faeces. To filter samples, we used a 330μm SaranTM mesh (Asahi Kasei, Japan). To reduce the amount of formaldehyde consumed in the process, we used saline instead of formaldehyde for all intermediate washing steps during concentration. For all centrifugation steps, we spun our samples at 3000 rpm for 5 minutes.

After processing, we again suspended the retained faecal pellet in SAF and stored it at room temperature until analysis. For parasitological examination, we used a volumetric method based on McMaster microscopy to estimate the intensity of parasitic infection in each individual through counting the number of eggs observed per gram of faecal sediment (EPG) (Modrý et al., 2018). We drew aliquots from the faecal suspension, kept homogeneous during reading using a magnetic stirrer, and viewed them in a McMaster chamber with a 10x objective lens. Parasite eggs were identified by their specific morphology and size (Modrý et al., 2018). We repeated this procedure five times per sample and used the mean count to calculate EPG from each sample based on the amount of sediment viewed per volume of suspension, with the following formula:

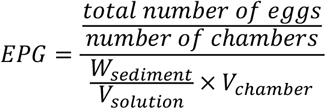

In the formula, total number of eggs reflects the number of eggs the examiner counted in all chambers. The number of chambers refers to the number of chambers examined in the sample (here, 5). W_sediment_ refers to the weight of the faecal pellet after filtration, and V_solution_ refers to the volume of solution that was used to dilute the sample (in this case, between 10ml and 50ml depending on the thickness of the faecal suspension, which significantly affects viewing). V_chamber_ refers to the volume of suspension contained in each counting chamber (0.15ml for a single McMaster chamber). EPG was then rounded to the nearest integer.

EPG is a standard index of parasite infection intensity. Although it is not always indicative of true worm infection intensity (Gillespie, 2006; Modrý et al., 2018), numerous studies have found significant correlations between EPG and the number of reproductive female worms infecting a given host (De Bont et al., 2002; Mohammed et al., 2016). The method is commonly used to measure domestic animal health and in wildlife parasitology, particularly in cases where parasite population estimation is needed but it is neither advisable nor possible to sample individuals via other - more invasive or destructive - means.

### Statistical Analyses

All statistical analyses were conducted using R v4.0.5 (R Development Core Team, 2021) with R studio (R Studio Team, 2021). We built a dataset that included one line for each parasite species in each faecal sample of each individual as the unit of analysis, i.e., three times (for the three parasite species) the number of faecal samples analysed. In addition, each sample was associated with an individual and all of its attributes, including its centrality measures (degree, strength, and eigenvector), Elo-rating, sex, and age, along with the sample’s collection date. EPG (rounded to the nearest integer) was used as the response variable in the statistical models described below. We also scaled and centred to 0 the centrality measures, age and Elo-rating.

#### Testing for associations between network centrality and parasite infection intensity

Considering that all parasites of interest are transmitted via contact with contaminated substrates, we decided to include EPG from different parasite species in one model as the response variable, accounting for variation among parasites with a control factor “parasite species”. In doing so, we accept the limitation that we are not testing for specific associations related to each parasite type, assuming instead that similar mechanisms may drive links between social behaviour and infection, should such links exist. We set parasite species identity as a fixed effect in the model to at least account for variation in EPG by species. We modelled rounded EPG as overdispersed count data. Parasites typically exhibit an aggregated distribution across their hosts that is best approximated by a negative binomial distribution (Crofton, 1971; Shaw et al., 1998; Poulin, 2007). We confirmed that the EPG distribution indeed followed a negative binomial distribution using the “fitdistr” function from the R package “MASS” (Venables and Ripley, 2002). We then built generalized linear mixed models with a negative binomial error structure using the “glmmTMB” package (Brooks et al., 2017).

The three centrality measures we investigated were related to each other (Spearman’s Correlation of degree and strength: r = 0.42, p = 0.002; degree and eigenvector: r = 0.47, p < 0.001; strength and eigenvector: r = 0.72, p < 0.001). Moreover, eigenvector centrality itself is derivative of degree and strength. Thus, including these metrics in the same model would violate the assumption of data independence, and the covariance between them could make the interpretability of coefficients challenging. Therefore, we constructed one model for each centrality measure. Age, sex, and Elo-ratings were set as control fixed effects as they are known to affect EPG but are not factors of interest here. EPG can also vary considerably over time and sample to sample within an individual (Wood et al., 2013). To account for such effects, we collected multiple faecal samples per individual and set the individual ID and date of sample collection as random effects in the models. The model residuals showed substantial heteroscedasticity and overdispersion, so we added a zero-inflation term (intercept-only) to account for the substantial number of zeros in the EPG data (124 instances or 24% of the 517 total data points). The zero-inflated models showed better residual distributions and significantly lower AIC compared with standard negative binomial models (degree: 7268.8<7727.0; strength: 7254.5<7718.0; eigenvector centrality: 7260.7<7722.1). We also assessed whether other assumptions of the models were met, such as lack of multicollinearity (variance inflation factor < 3) and overly influential cases (Cook’s distance < 4/N). Lastly, we conducted likelihood ratio tests (LRT) to evaluate the statistical strength of our models against an informed null model containing only the control factors age, sex, Elo-rating, parasite species, and random effects.

Note that we did not carry out any permutation or randomisation procedure typically associated with non-independent data such as social network centrality measures to account for autocorrelation induced by the nature of the data (Hart et al., 2021; Weiss et al., 2021; Farine and Carter, 2022). In this study, centrality metrics were treated as predictor variables, not as response variables. The statistical issue of non-independence arises when network measures are the response variable because it is important that residuals are independent, which relates to links between the response and the predictor variable(s). In this study, we include network centrality metrics as predictors like other individual traits such as rank or age (which can also be considered inter-dependent; yet, we do not habitually control for it). Here, mixed models should be able to account for dependencies in the data if they are well specified, as we believe they are.

#### Testing the effect of knockout simulations on the relationship between centrality and parasite infection intensity

To have a point of comparison with previous studies and to gain insight into the influence of only examining certain age-sex classes on the relationship between centrality and infection intensity, we ran two other analyses. We first constructed partial social networks including only (1) adult females (N=20, or 38.5% of the group), which mimics what was done in previous studies, or (2) juveniles (N=28, or 53.8% of the group). We calculated Spearman’s correlation coefficients to compare between the network metrics from original and partial networks. Then, we built models with the same structure as those described above but with network metrics calculated from partial networks. These models went through all the checks as previously described.

Removing individuals from the network itself might already affect social network structure and thus change the observed result. In a second step, therefore, we conducted randomized knock-out simulations: we randomly removed 5%, 10%, 25%, and 50% of individuals from the network, re-calculated the network metrics, and re-integrated them into the models for calculation over 1000 iterations. After each simulation, the model parameters of interest (here estimates and confidence intervals) were saved and the results were compared with the results from the original models. We chose those percentages because they represent biologically relevant cases of missing individuals during data collection or within a study design, like solitary roaming males or peripheral females (e.g., <10% of the group), or when taking into account only certain age-sex classes, here adult females or juveniles, effectively removing everyone else (e.g., >25% of the group, depending on group composition and demography).

Knock-out simulations not only disrupt the network structure and change the centrality metrics calculated, but they also reduce the number of individuals included in the model, reducing sample size and possibly statistical power. We therefore further analysed the effect of reducing the number of individuals in the networks using the following approach. We first sub-sampled the complete network as before (i.e. adult female-only data or juvenile-only data), but instead of re-constructing the network and re-calculating metrics based on this reduced set of individuals, we ran the statistical models using the centrality metrics for those individuals from the original whole-group network. We then re-ran the same randomized knock-out procedure, but this time retaining the centrality metrics from the original whole-group network. In this way, we are able to determine to what extent any differences between results with subsampled networks and with the original network stem from real changes in network structure versus issues caused by reduced statistical power (i.e. smaller sample size). If the models’ centrality estimates without network metric recalculation are found to be of the same magnitude as the models’ centrality estimates with network recalculation, then results would be independent of the way the centrality measures are calculated and would have little to do with missing interactions, which might be rather indicative of a power issue (fewer data points).

## Results

### Social network metrics

The whole-group network includes 52 nodes with 480 connections between nodes, and is based on 14,420 proximity interactions. In the whole-group network, individuals on average had 14 (± 5) social partners (degree centrality), an average proximity strength centrality of 0.44 ± 0.21 and an average eigenvector centrality of 0.32 ± 0.29 (Table 2). The adult female-only network included 20 nodes with 87 connections between nodes, and was based on 2,403 proximity interactions. The juvenile-only network included 24 nodes with 139 connections, and was based on 2,316 proximity interactions (see Table 1 for details).

**Table 1:** Summary of network metrics from whole-group or partial networks in this study

| | Whole-group<br>(mean±SD) | Adult females<br>(whole-group<br>network)<br>(mean±SD) | Juveniles<br>(whole-group<br>network)<br>(mean±SD) | Adult females<br>(adult female-only network)<br>(mean±SD) | Juveniles<br>(juvenile-only<br>network)<br>(mean±SD) |
| --- | --- | --- | --- | --- | --- |
| Degree | 13.80±5.48 | 13.75±4.22 | 15.33±4.93 | 6.20±2.76 | 8.83±2.79 |
| Strength | 0.44±0.21 | 0.55±0.20 | 0.43±0.16 | 0.21±0.11 | 0.15±0.09 |
| Eigenvector | 0.32±0.29 | 0.39±0.32 | 0.33±0.29 | 0.33±0.32 | 0.25±0.29 |

**Table 2:**
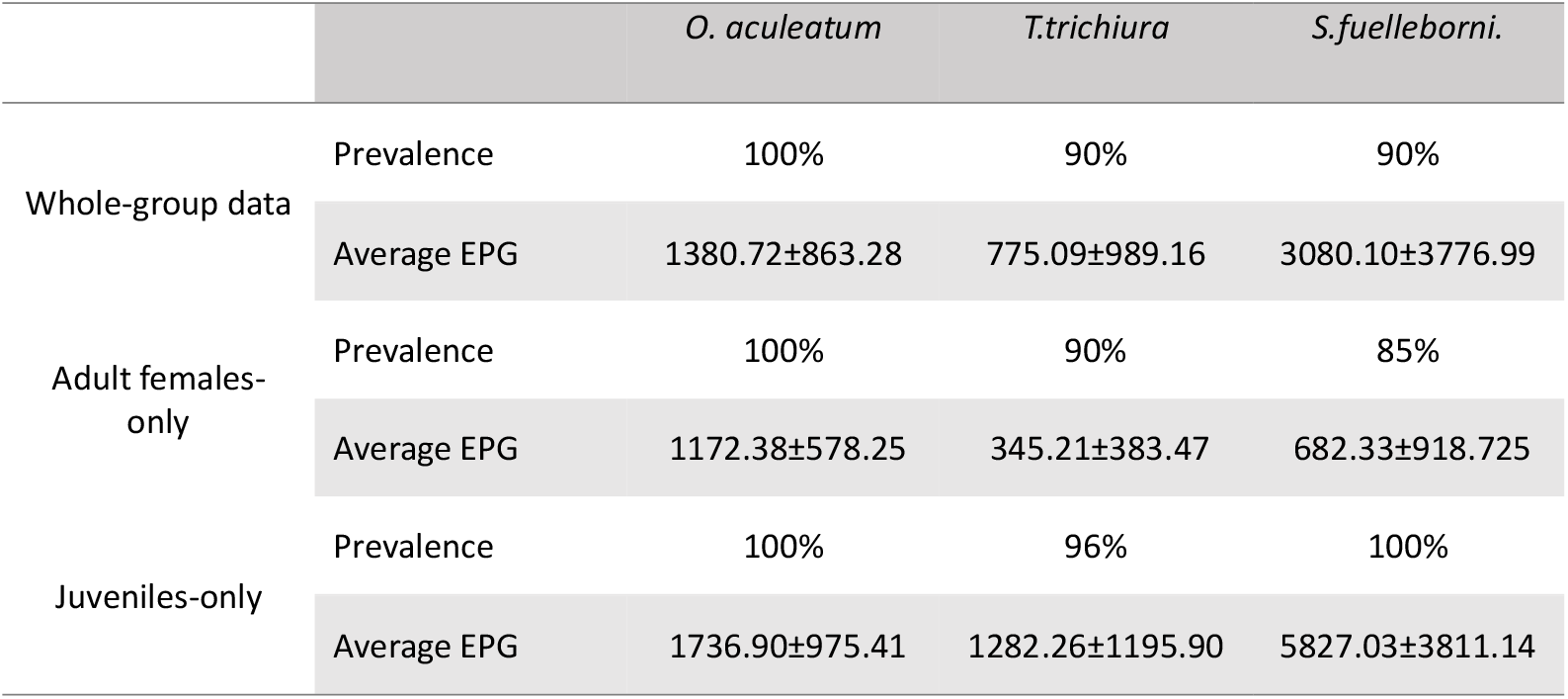
Summary of prevalence and infection intensity (EPG) of three geohelminths examined in this study.

|  |  | *O. aculeatum* | *T. trichiura* | *S. fuelleborni* |
| --- | --- | --- | --- | --- |
| Whole-group data | Prevalence | 100% | 90% | 90% |
| | Average EPG | 1380.72±863.28 | 775.09±989.16 | 3080.10±3776.99 |
| Adult females-only | Prevalence | 100% | 90% | 85% |
| | Average EPG | 1172.38±578.25 | 345.21±383.47 | 682.33±918.725 |
| Juveniles-only | Prevalence | 100% | 96% | 100% |
| | Average EPG | 1736.90±975.41 | 1282.26±1195.90 | 5827.03±3811.14 |

### Parasitological results

We observed all three of the expected geohelminths infecting our subjects during the study period: *Oesophagostomum aculeatum, Strongyloides fuelleborni, Trichuris trichiura*. Most individuals were infected by all parasites (see Table 2 for details).

### Network-infection relationship

#### Whole-group networks

Likelihood ratio tests demonstrated that the models with strength and eigenvector centrality, but not with degree centrality, as fixed effects had significantly higher explanatory value than their respective informed null models (Table 3). Strength and eigenvector centralities were positively related to infection intensity, such that more central individuals had higher geohelminth EPG (Table 3). In addition, age was negatively related to infection, with younger individuals showing higher EPG. Sex and dominance rank were not related to infection intensity (Figure 2).

**Table 3:** Results of GLMMs testing for variation in geohelminth infection intensity (EPG) among a whole group of Japanese macaques on Koshima Island. LRT refers to likelihood ratio test, here shown with Chi-square χ2 tests, degrees of freedom (d.f.) and p-values (p). Estimates (β) are given with standard errors (SE) and p-values (p). Variance is given with standard deviation (SD). Results in bold are statistically significant (p < 0.05). For clarity, results from models that failed to outperform their null counterparts are presented in full in the table SI-4.

| *Centrality metric of focus* | *Degree* | *Strength* | *Eigenvector* |
| --- | --- | --- | --- |
| *LRT against null*<br>($\chi^2$; d.f.; p) | 1.940; 2;<br>0.379 | 16.491; 2;<br><0.001 | 10.238; 2; 0.006 |
| Fixed effects | *Estimate $\beta \pm$ SE*<br>[p] |  |  |
| Intercept | / | 6.88 ± 0.12<br>[<0.001] | 6.89 ± 0.13<br>[<0.001] |
| Centrality metric | / | **0.29 ± 0.09**<br>[0.003] | **0.21 ± 0.09**<br>[0.025] |
| Age | / | **-0.53 ± 0.13**<br>[<0.001] | **-0.45 ± 0.13**<br>[<0.001] |
| Sex (male)* | / | 0.34 ± 0.21<br>[0.110] | 0.29 ± 0.22<br>[0.199] |
| Elo-rating | / | -0.16 ± 0.11<br>[0.152] | -0.13 ± 0.12<br>[0.286] |
| Species (Stro)** | / | **0.65 ± 0.13**<br>[<0.001] | **0.65 ± 0.13**<br>[<0.001] |
| Species (Tri)*** | / | **-0.67 ± 0.12**<br>[<0.001] | **-0.67 ± 0.12**<br>[<0.001] |
| Random effects | *Variance* |  |  |
| Collection Date | / | 0.02 | 0.02 |
| Individual ID | / | 0.24 | 0.26 |

\* Sex (male) shows model coefficients for males compared with the reference value, females
\*\* Species (Stro) shows model coefficients for the parasite species *S. fuelleborni* compared with the reference value, *O. aculeatum*
\*\*\* Species (Tri) shows model coefficients for the parasite species *T. trichiura* compared with the reference value, *O. aculeatum*

**Figure 1:**
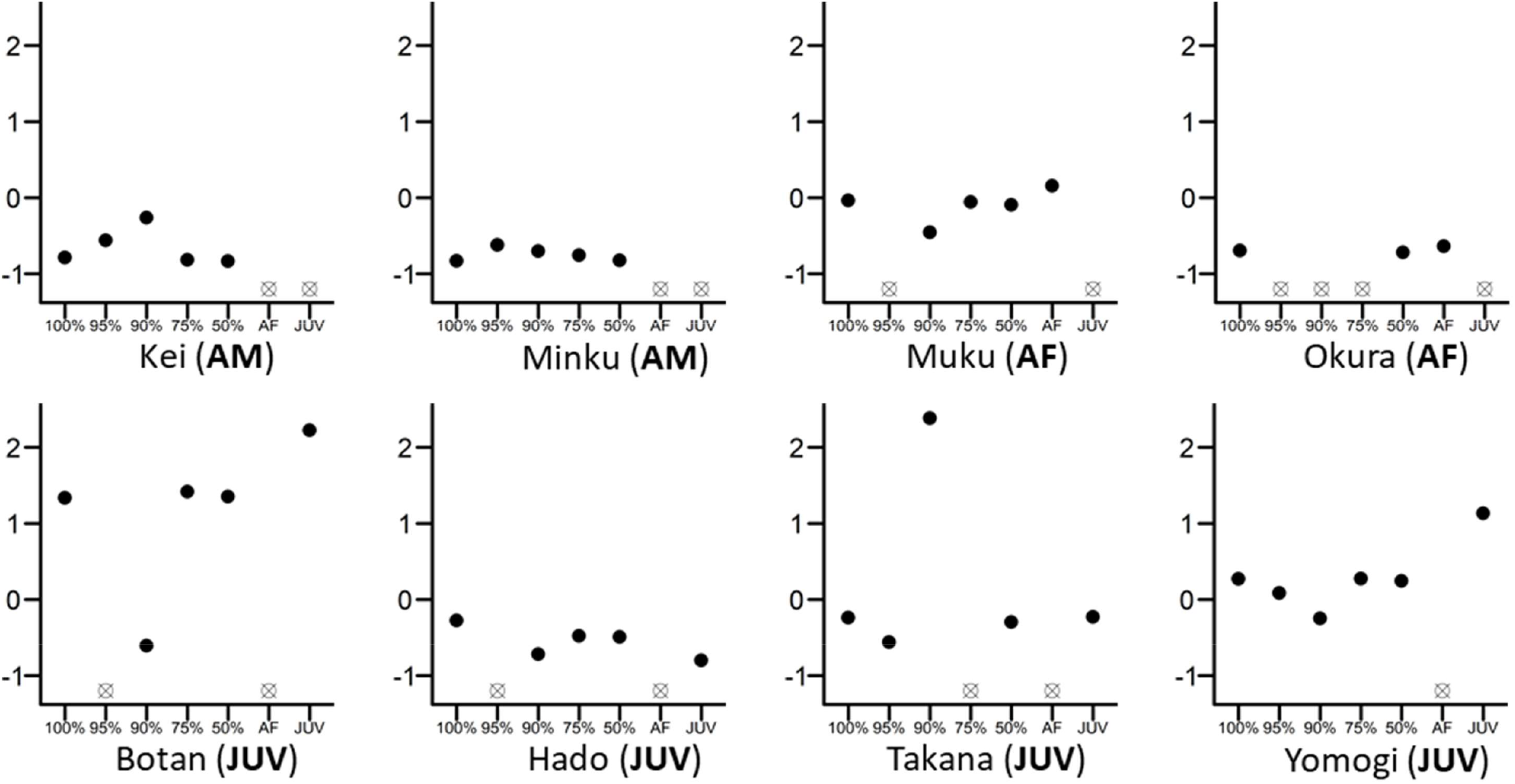
Differences in the eigenvector centralities of eight randomly chosen individuals from the whole group. The 100% mark on the x-axis reflects the whole-group network, while 95%, 90%, 75%, 50% are values retained from networks where 5%, 10%, 25% or 50% of individuals were randomly removed. Individual figures are marked as AM (adult male), AF (adult female) or JUV (juvenile). All eigenvector centrality values were scaled and centred to 0 within their own network for direct comparison. Empty crossed dots mean that the individual was either removed randomly from that network or that the individual does not belong in this particular network; for example, the individual is an adult female and cannot figure into the juvenile-only network (e.g. Okura (AF)).

**Figure 2:**
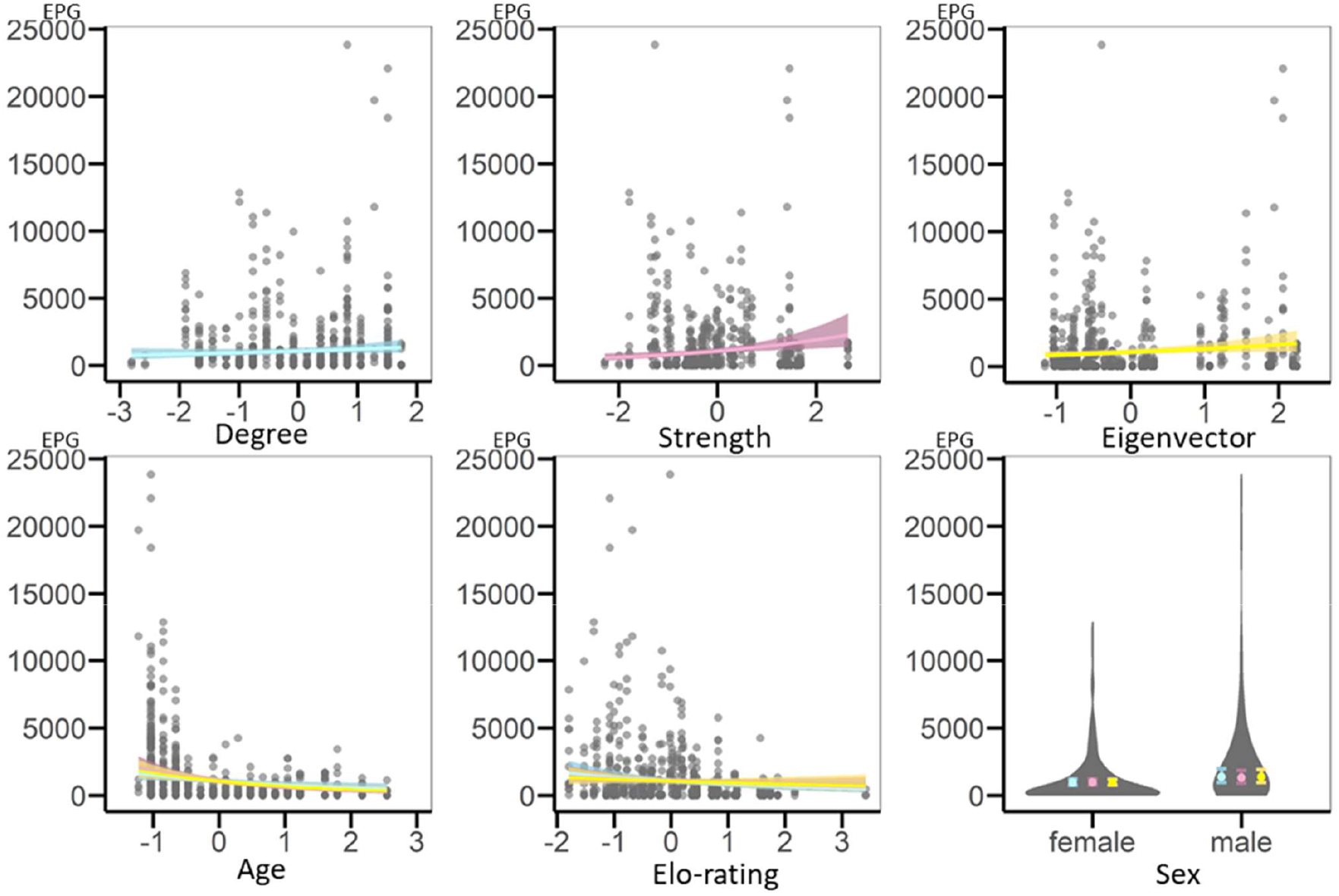
Regression plots representing results of GLMMs testing for variation in geohelminth infection intensity (EPG) based on the whole-group network. On the x-axis are the scaled variables, on the y-axis is infection intensity (EPG). Each circle is a data point. Each network metric is included in its own model and is represented by its own colour, blue for the model with degree, pink for the model with strength and yellow for the model with eigenvector centrality. The regression plots representing shared fixed effects (age and Elo-rating) show lines from each of the models in the respective different colours. Note that results are very similar and thus lines overlap. The plot for sex is a violin plot representing the spread of the data (violins) and the estimates with standard errors (filled circles and error bars) shown in each model’s respective colour.

#### Female- and juvenile-only networks

When comparing measures calculated from partial versus whole-group networks, in both adult female-only and juvenile-only networks, most measures derived from partial-networks were strongly to moderately correlated with the original whole-group network measures. Only strength derived from the adult female-only network was not correlated with strength derived from the original whole-group network (Spearman’s rank correlation; adult females: r_degree_ = 0.71, p < 0.001, r_strength_ = 0.07, p = 0.772, r_eigenvector_ = 0.76, p < 0.001; juveniles: r_degree_ = 0.87, p < 0.001, r_strength_ = 0.80, p < 0.001, r_eigenvector_ = 0.69, p < 0.001).

Using data from adult females only, likelihood ratio tests demonstrated that the models with strength, but not those with eigenvector centrality or degree centrality as main predictors, had higher explanatory value than their respective informed null models (Table 4). Neither age nor rank was related to nematode infection intensity (Table 4, Figure SI-2). Using data from juveniles only, likelihood ratio tests demonstrated that none of the models had a higher explanatory value than their respective informed null models (Table 5, Figure SI-3). The same models after data reduction (i.e. with centrality measures derived from the whole-group network) showed similar results (in terms of estimate values in the model) to those after network reduction (i.e. based on recalculated measures from sub-sampled networks), except for juvenile-only networks, which gave much lower model estimates overall (see Table SI-7, Table SI-8). In the case of adult females, it seemed that regardless of how the network was built (i.e. with more or fewer interactions/connections), results stayed more or less the same. In the case of juveniles, however, ignoring the fact that models still did not differ from their null counterparts, model estimates were much lower with more interactions/connections included. These results suggest that, depending on the (sub-)population sampled, either a loss of statistical power with decreasing sample size or a change in the network itself can lead to divergent results concerning associations between centrality and infection.

**Table 4:** Results of GLMMs testing for variation in geohelminth infection intensity (EPG) among adult female Japanese macaques on Koshima Island. LRT refers to the likelihood ratio test, here shown with Chi-square χ2 tests, degrees of freedom (d.f.) and p-values (p). Estimates (β) are given with standard errors (SE) and p-values (p). Results in bold are statistically significant (p < 0.05). For clarity, results from models that failed to outperform their null counterparts are presented in full in table SI-5).

| *Centrality metric of focus* | *Degree* | *Strength* | *Eigenvector* |
| --- | --- | --- | --- |
| *LRT against null*<br>($\chi^2$; d.f.; p) | 3.616; 2; 0.164 | **7.424; 2; 0.024** | 4.214; 2; 0.122 |
| Fixed effects | *Estimate $\beta \pm SE$*<br>*[p]* |  |  |
| Intercept | / | 6.76 ± 0.13<br>[<0.001] | / |
| Centrality metric | / | **0.32 ± 0.12**<br>**[0.006]** | / |
| Age | / | -0.09 ± 0.13<br>[0.477] | / |
| Elo-rating | / | -0.06 ± 0.13<br>[0.622] | / |
| Species (Stro)* | / | **-1.00 ± 0.29**<br>**[<0.001]** | / |
| Species (Tri)** | / | **-1.16 ± 0.18**<br>**[<0.001]** | / |
| Random effects | *Variance* |  |  |
| Collection Date | / | 0.00 | / |
| Individual ID | / | 0.15 | / |

\* Species (Stro) shows model coefficients for the parasite species *S. fuelleborni* compared with the reference value, *O. aculeatum*
\*\* Species (Tri) shows model coefficients for the parasite species *T. trichiura* compared with the reference value, *O. aculeatum*

**Table 5:** Results of GLMMs testing for variation in geohelminth infection intensity (EPG) among juvenile Japanese macaques on Koshima Island. LRT refers to the likelihood ratio test, here shown with Chi-square χ2 tests, degrees of freedom (d.f.) and p-values (p). For clarity, results from models that failed to outperform their null counterparts are presented in full in table SI-6).

| *Centrality metric of focus* | *Degree* | *Strength* | *Eigenvector* |
| --- | --- | --- | --- |
| *LRT against null*<br>($\chi^2$; d.f.; p) | 0.120; 2; 0.942 | 2.491; 2; 0.288 | 1.643; 2; 0.440 |

#### Random partial networks

Random removals of individuals comprising 5, 10, 25, and 50% of the group, following which networks were reconstructed with remaining individuals, all led rapidly to a decrease in the strength of the observed relationship between infection and centrality (except for degree, in which case no relationships were observed in both original network and partial networks) (Table 6). In other words, in these models, estimates describing that observed relationship approached zero and did not reach statistical significance (Figure SI-9). Moreover, these results do not appear to be caused by a loss of statistical power, because the majority of results from models in which knockouts were conducted but centrality measures were retained from the whole-group network were much closer to the original results obtained with all individuals present (Table SI-10, Figure SI-11).

**Table 6:** Results of 1000 random removals of individuals (% given in the first column) run with observed network data and used to (re)model the relationship between network centrality and infection intensity. The heading “Number of model results equivalent to observed” indicates the number of simulation model results out of 1000 simulations that gave an equivalent result to models based on the whole-group network, i.e. for strength and eigenvector, positive estimates and 95% confidence intervals not overlapping zero, or for degree, a near-zero estimate with 95% confidence intervals overlapping zero.

| *Centrality measure* | Percentage of individuals removed | Number of model results equivalent to observed | Number of significant model results in the opposite direction |
| --- | --- | --- | --- |
| Degree | 5% | 98.0% | / |
|  | 10% | 96.6% | / |
|  | 25% | 95.4% | / |
|  | 50% | 95.9% | / |
| Strength | 5% | 3.2% | 0.7% |
|  | 10% | 4.2% | 1.5% |
|  | 25% | 3.9% | 4.0% |
|  | 50% | 4.7% | 5.1% |
| Eigenvector | 5% | 12.6% | 0.7% |
|  | 10% | 7.1% | 1.2% |
|  | 25% | 3.8% | 3.8% |
|  | 50% | 5.3% | 5.2% |

## Discussion

Empirically and theoretically, increased sociality is oftentimes linked to increased parasitism, although this relationship is not always linear (Freeland, 1976; Altizer et al., 2003; Griffin and Nunn, 2012; Godfrey, 2013; Nunn et al., 2015, 2015; Romano et al., 2016; Rushmore et al., 2017; White et al., 2017; Briard and Ezenwa, 2021; Lucatelli et al., 2021). Our study found positive correlations between geohelminth infection intensity and two out of three social network centrality measures examined in a Japanese macaque group. Specifically, the frequency with which individuals were in close proximity to one another (network strength), as well as their further social integration (eigenvector centrality), but not the number of individuals with which they shared proximity (degree centrality), were positively associated with infection intensity of geohelminths. These results partially align with the results of previous work in a different population of Japanese macaques (MacIntosh et al., 2012), with both investigations suggesting that eigenvector centrality, whether in grooming or proximity networks, is associated with nematode parasitism in Japanese macaques.

Our results also align with a recent meta-analysis of studies investigating parasites in host social networks, wherein strength and eigenvector centrality showed stronger associations with parasitism than did any of betweenness, closeness, or degree centrality (Briard and Ezenwa, 2021). It therefore seems important to consider what these various centrality measures represent in terms of host socio-biology and in terms of social or ecological processes. Exposure and transmission risk of fast and directly spread parasites such as lice or certain viruses and bacteria might depend simply on the number of associations individuals have (degree), whereas slower-spreading/developing parasites or those requiring additional developmental stages in the environment may be more dependent on the actual time hosts spend in contact or in spatial proximity with one another (strength), or the overall degree of social integration they have (eigenvector centrality) (Silk et al., 2017; Briard and Ezenwa, 2021; Lucatelli et al., 2021). By whatever mechanism, our results support those of review studies and meta-analyses showing that, across many different sociality measures and parasites with different life histories and modes of transmission, the relationship between sociality and parasitism is generally positive (Altizer et al., 2003; Rushmore et al., 2017; Briard and Ezenwa, 2021; Lucatelli et al., 2021).

However, we also demonstrated that the choice of animals within a study group (i.e., sampling bias) can impact the relationship that is observed between network centrality and parasitism, a fact that is often neglected or ignored in wildlife studies. Indeed, numerous studies have focused on specific age-sex classes rather than the whole social group (MacIntosh et al., 2012; Duboscq et al., 2016; Romano et al., 2016; Webber et al., 2016; Hamilton et al., 2020; Roberts and Roberts, 2020; Sandel et al., 2021). In our study, considering partial networks – e.g., those including only a subset of the group like adult females or juveniles – yielded divergent results compared to those analyses that included the whole-group network. Statistical relationships originally observed between network centrality and infection intensity weakened considerably, except for network strength in the adult female model, although the direction of the effect remained positive in all cases for both strength and eigenvector centrality. Thus, sampling bias can lead to misdirected conclusions about the relationships between sociality and parasitism, and potentially any other network process.

From an epidemiological perspective, every individual can be responsible for the transmission of a pathogen, so neglecting them in the analysis may reduce the reliability of the data. In our study, as in previous work with Japanese macaques (MacIntosh et al., 2010), descriptive statistics show that juveniles exhibited higher geohelminth infection intensities than adults. Furthermore, within the juvenile class, younger individuals and those that were male shed more parasite eggs than did older individuals or those that were female (Table SI-5). Although these models did not reach statistical significance, these results concur with the general findings that young animals and males are disproportionately infected by parasites (Khan et al., 2010; Hinney et al., 2011; Habig and Archie, 2015). Hypotheses explaining this difference include age/sex-associated hormonal profiles (Roberts et al., 2001), variable levels of immunity (Roberts et al., 2001; Fallon et al., 2003; East et al., 2008; Ebersole et al., 2008), different probabilities of exposure to contaminated items (Sarabian, Belais, et al., 2018; Sarabian, Curtis, et al., 2018), and differing social positions within the group (Grassi, 2002; Cords et al., 2010; Liao et al., 2018). Social positions of juveniles in particular are still under development and are thus less stable than those of adults (Kulik et al., 2015). Therefore, social network centrality estimates based on their behaviour might be less representative of the actual relationships, or simply contain far more noise, and therefore fail to capture network-infection correlations. Considering such age and sex biases in infection, along with the variable activity patterns (in terms of social interactions and ranging behaviour) performed by juveniles and males, neglecting them as study subjects may have important consequences for the conclusions reached about links between sociality and infection. Moreover, because individuals form connected nodes in networks and are thus not independent of one another, excluding them from the network fundamentally changes its overall structure (Fedurek and Lehmann, 2017). Indeed, missing nodes can dramatically change network structure and affect the metrics calculated (Kossinets, 2006; Smith and Moody, 2013; Silk et al., 2015; Smith et al., 2017; Silk, 2018). Such effects may percolate through the processes linked to network structure, such as the transmission of information or pathogens (Godfrey et al., 2010).

That said, in our study at least, most of the network metrics calculated from both the adult female-only network and the juvenile-only network were rather well correlated with those calculated from the whole-group network. Among these correlations, degree and strength from the juvenile-only network were closer to those from the whole-group network than were the same measures from the adult female-only network. And, interestingly, strength from the adult female network was not correlated with the whole-group metric. In contrast, eigenvector centralities from the adult female-only network were closer to those in the whole-group network than were eigenvector centralities from the juvenile-only network. A possibility would be that each juvenile is likely to interact with a small number of adults, i.e., adult female kin, and most of their contacts are between juveniles. Therefore, the inclusion or exclusion of adult individuals in a juvenile network may have comparatively less influence on their degree or strength centrality than the reverse. In contrast, each adult female may have multiple offspring, and this number can vary across individuals, with the potential to significantly change the number of contacts a female has, and the frequency with which they interact. With regards to eigenvector centrality, this measure represents the integration of an individual within its whole network of connections. Since females are known to form the core of Japanese macaque groups (Yamagiwa and Hill, 1998), it may be that their patterns of interaction dominate the emergent social network structure, such that properties of female networks linked to social integration (as in eigenvector centrality) do not change as much with the inclusion/exclusion of other subgroups.

In stark contrast to these targeted knockouts, our random knockouts quickly erased any relationship found in the original network between infection and centrality, for all metrics observed. This was evidenced both by the complete absence of statistical significance in the knockout models, which was rapidly lost even at just 5% random reduction in individuals in the network, and by the magnitude of the model estimates, which quickly approached zero at just 5% random network reduction. Random removal of individuals in a network then, much more so than targeted removal of subsets, might cause greater disruption in network structure, at least as far as it concerns processes occurring on networks, like infection. It has been shown that targeted removal of central individuals from a network has greater effects on resultant network measures than random removal (Smith and Moody, 2013; Smith et al., 2017). We did not test the effect of removing specific individuals in our study, so cannot draw comparison here, but that is a topic worthy of future consideration. For example, the chance of removing at least one adult female from the studied network would be 77% when removing 5% of individuals from the group at random. As adult females form the core of Japanese macaque groups (Yamagiwa and Hill, 1988), removing only a few of them might have dramatic and cascading effects on the network structure overall. Furthermore, it has been shown that, as the proportion of individuals excluded from a network increases, the probability that one or more central nodes are excluded also increases, leading to stronger effects on the resultant metrics (Silk et al., 2015; Smith et al., 2017; Silk, 2018). However, since our random knockouts resulted in a complete loss of the infection-centrality link at just 5% removed, this is unlikely to be a key factor explaining our results.

Now, in real world cases, missing individuals from a network is rarely a random process, especially if significant study or observation effort is made over extended time periods. Instead, there are usually systematic reasons for missing certain individuals, for example those that are difficult to observe, such as roaming adult males or young, unrecognisable juveniles. This is one reason why we focused on removing specific sex/age classes from our data set. It is notable that, in our study, complete age/sex subsets preserved the infection-centrality link far better than the random knockouts did. This might suggest that, while the full network is always preferred, including complete subsets might retain much more of the information present in the original network than would taking a random sample of individuals. It will be interesting to examine these effects in additional species of hosts and parasites, especially those with different network structures or modes of transmission, to further disentangle the impact of social dynamics and sampling on network processes like parasite transmission. Ultimately, whether individuals are omitted intentionally or accidentally, our results highlight the potential impacts of partial networks on the conclusions drawn.

One final point to consider is the potential reasons behind the different results obtained in the knock-out simulations: the act of removing nodes itself, i.e., a data reduction issue leading to statistical power issues, or the actual disruption in the network structure caused by removing nodes and their perceived relationships. We posited that if the model centrality estimates with data reduction (without network metric recalculation) were of the same magnitude as the model centrality estimates with network restructuring (and recalculation), then results can be seen as independent of the way the measures are calculated. If so then, results have little to do with missing interactions which might be rather indicative of a power issue (less data points). If we look at the change in centrality estimates from all models regardless of whether they outperformed their null counterparts (Table SI-5 & Table SI-7; Table SI-6 & Table SI-8), we obtained contrasting outcomes: in adult females, results are unchanged for degree and strength but for eigenvector centrality, model estimates with data reduction are lower than estimates with network reduction. In juveniles, model estimates with data reduction are several-fold lower than estimates with network reduction. Therefore, we cannot distinguish whether the divergent results were generally caused by disruption of the network or because removal of specific set of data points reduced statistical power of the model when including only specific age-sex classes. If we look at the change in centrality estimates in the random knockouts, however, the results show clear differences: estimates from simulations with network restructuring centred on zero, while estimates from simulations with data reduction centred on original (expected) results (Figure SI-9, Figure SI-11). Thus, disruption of the network caused by missing nodes is the main reason why divergent results were obtained after random removal of individuals.

Despite our results and the conclusions drawn from them, we do not intend to devalue the results of previous studies solely on these grounds. Partial networks can still provide robust network metrics, providing that there is little imbalance as to who gets excluded (Borgatti et al., 2006; Silk et al., 2015). In our study, we focused only on excluding broad categories of individuals or used pseudo-random knockouts, and we did not control for the specific social characteristics of the removed individuals. It is further possible that excluding particular individuals – super-spreaders, keystone individuals, policing individuals, alpha individuals – would have a much larger effect on the processes that occur on social networks than would excluding individuals with a less influential role. For instance, knocking out policers in a social group of pigtailed macaques greatly destabilized the structure of several social networks, setting the stage for reduced group cohesion (Flack et al., 2006). This is also reminiscent of work investigating the effect of targeting specific individuals for health interventions or information transmission: depending on which individual is targeted, information has a greater or lesser likelihood of spreading in the community (e.g. Kim et al., 2015). Similar work has looked at the spread of an infectious disease or a parasite depending on which individual gets it first, and what position they occupy in the social network (e.g. Romano et al., 2016). So, knowing the characteristics of individuals that play important roles in the sociality-parasitism relationship can help practitioners develop strategies, for example, for disease control (Rushmore et al., 2014; Romano et al., 2016) and captive population management (McCowan et al., 2008). The key, however, is to ensure that the assumptions made are the result of careful consideration of the completeness and thus representativeness of the data at hand.

## Acknowledgements

We are grateful to the Cooperative Research Program of Kyoto University’s Wildlife Research Center and the city of Kushima’s Agency for Cultural Affairs for permitting this research. We thank Christof Neumann for statistical and programming support, accepting that all analytical decisions – and outstanding errors – are ours alone. We thank Dr. Zhang Peng for supporting this research and collaboration through Sun Yat-Sen University. Version 6 of this preprint has been peer-reviewed and recommended by Peer Community In Network Science (https://doi.org/10.24072/pci.networksci.100005).

## Data and scripts availability

Data are available online: https://doi.org/10.5281/zenodo.6825009

## Supplementary material

Supplementary material are available online: https://www.biorxiv.org/content/10.1101/2021.06.07.447302v5.supplementary-material.

