## Supplementary information for "Linking parasitism to network centrality and the impact of sampling bias in its interpretation"

Table SI-1. The first rows of the data set used in the analyses. NO = identity of faecal sample; ID = identity of individual; CD = date of faecal sample collection; D = degree centrality; S = strength centrality; EV = eigenvector centrality; A = age of the individual; AG = age group of individual; EL = Elo-rating of individual; FOA = total amount of observation scans per individual; FSA = total number of faecal samples collected per individual; Species = parasite species included. Note that these data are raw data, not yet scaled and centred. The complete data set is available from Git repository: https://github.com/XuZhihong-cn/Network_Centrality_analysis_and_KO-simulation

| *NO* | *ID* | *CD* | *D* | *S* | *EV* | *A* | *AG* | *SEX* | *EL* | *FOA* | *FSA* | *EPG* | *Species* |
|---|---|---|---|---|---|---|---|---|---|---|---|---|---|
| 2151 | beni | 14 | 18 | 0.385126 | 0.292249 | 8 | adult | female | 569 | 439 | 4 | 263.4468 | Oes |
| 2187 | beni | 21 | 18 | 0.385126 | 0.292249 | 8 | adult | female | 569 | 439 | 4 | 698.6028 | Oes |
| 2194 | beni | 23 | 18 | 0.385126 | 0.292249 | 8 | adult | female | 569 | 439 | 4 | 399.5006 | Oes |
| 2232 | beni | 29 | 18 | 0.385126 | 0.292249 | 8 | adult | female | 569 | 439 | 4 | 1674.419 | Oes |
| 2128 | kanna | 9 | 9 | 0.526477 | 0.042982 | 13 | adult | female | 763 | 487 | 3 | 2014.388 | Oes |

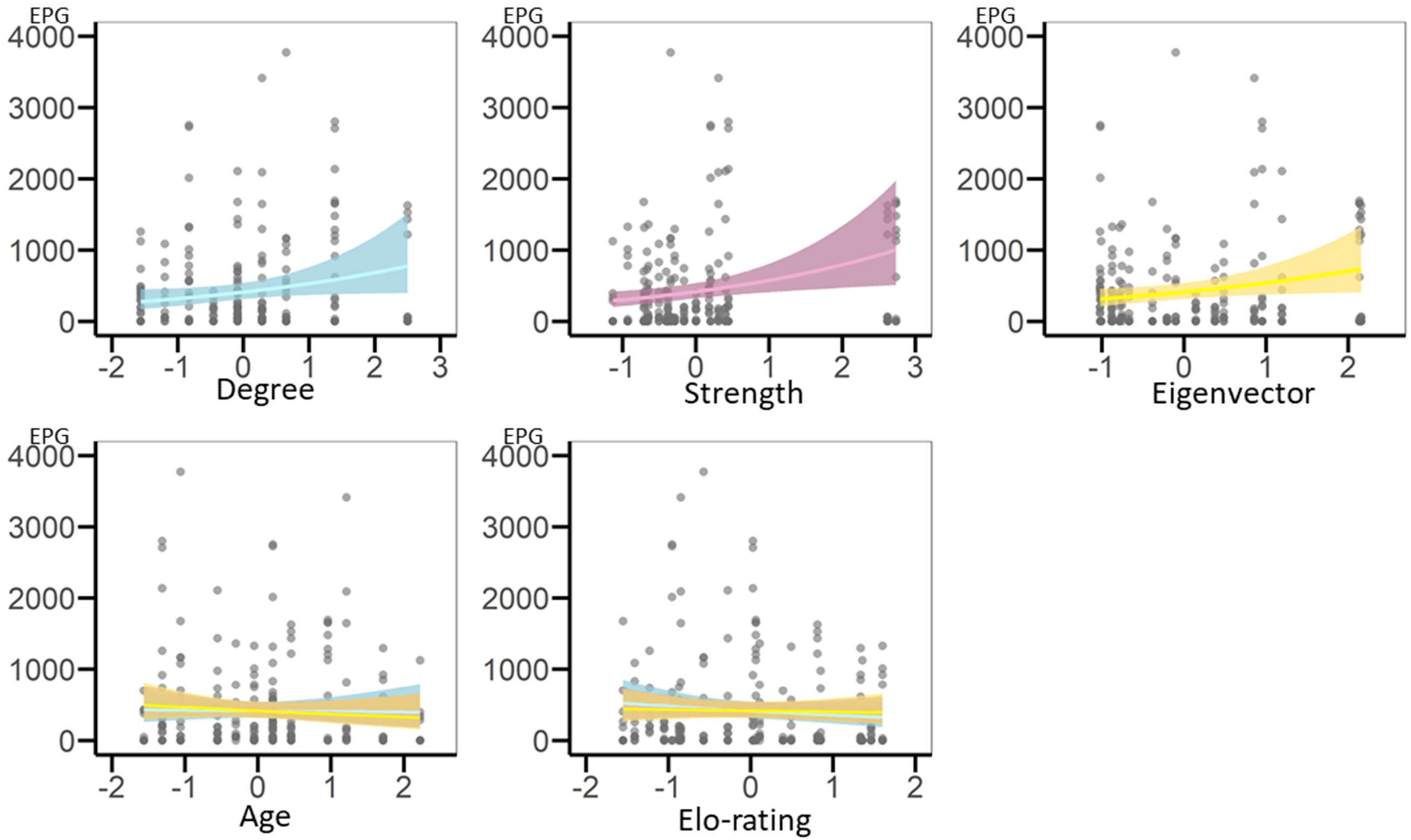


Figure SI-2: Regression plots representing results of GLMMs based on the **adult-female only network**. On the x-axis are the scaled variables, on the y-axis is infection intensity (EPG). Each circle is a data point. Each network metric is included in its own model and is represented by its own colour, blue for the model with degree, pink for the model with strength and yellow for the model with eigenvector centrality. The regression plots representing shared fixed effects (age and Elo-rating) show lines from each of the models in the respective different colours. Note that results are very similar and thus lines overlap.

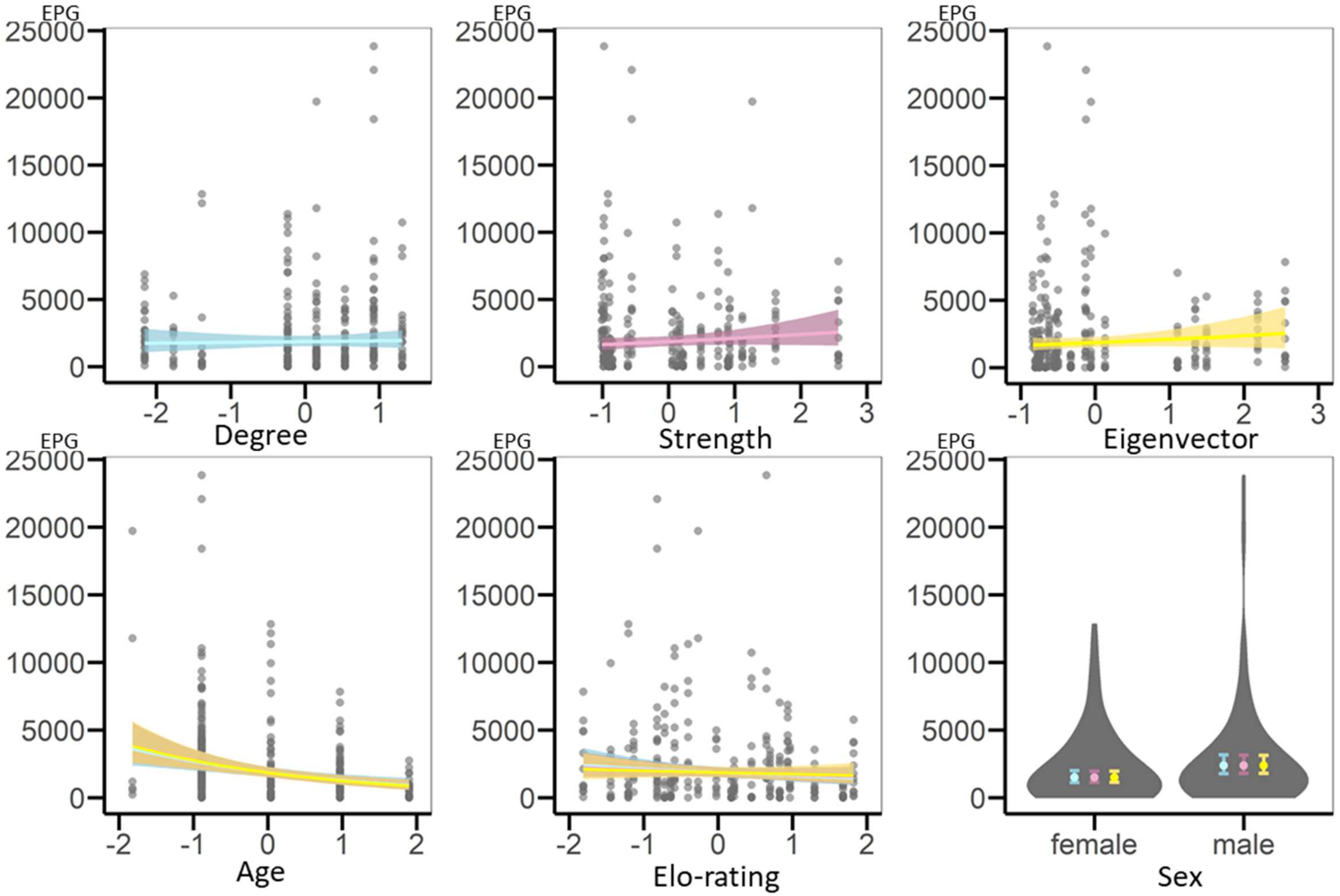


Figure SI-3: Regression plots representing results of GLMMs based on the **juvenile-only network.** On the x-axis are the scaled variables, on the y-axis is infection intensity (EPG). Each circle is a data point. Each network metric is included in its own model and is represented by its own colour, blue for the model with degree, pink for the model with strength and yellow for the model with eigenvector centrality. The regression plots representing shared fixed effects (age and Elo-rating) show lines from each of the models in the respective different colours. Note that results are very similar and thus lines overlap. The plot for sex is a violin plot representing the spread of the data (violins) and the estimates with standard errors (filled circles and error bars) shown in each model's respective colour.

Table SI-4. Results of GLMMs testing for variation in geohelminths infection intensity (EPG) among a **whole group** of Japanese macaques on Koshima island. LRT refers to likelihood ratio test, here shown with Chi-square $\chi^2$ tests, degrees of freedom (d.f.) and p-value (p). Estimates (β) are given with standard errors (SE) and p-value (p). Statistically significant ($p < 0.05$) results from models that outperform the null model are set in bold text.

| *Centrality metric of focus* | *Degree* | *Strength* | *Eigenvector* | *Null* |
|---|---|---|---|---|
| *LRT against null* ($\chi^2$; d.f.; p) | 1.970; 2; 0.379 | 16.491; 2; <0.001 | 10.238; 2; 0.006 | / |
| Fixed effects | *Estimate β* ± SE [p] | | | |
| Intercept | 6.88 ± 0.13 [<0.001] | 6.88 ± 0.12 [<0.001] | 6.89 ± 0.13 [<0.001] | 6.89 ± 0.13 [<0.001] |
| Centrality metric | 0.12 ± 0.09 [0.165] | **0.29 ± 0.09 [0.003]** | **0.21 ± 0.09 [0.025]** | / |
| Age | -0.33 ± 0.13 [0.009] | **-0.53 ± 0.13 [<0.001]** | **-0.45 ± 0.13 [<0.001]** | **-0.37 ± 0.13 [<0.001]** |
| Sex (male)* | 0.35 ± 0.23 [0.126] | 0.34 ± 0.21 [0.110] | 0.29 ± 0.22 [0.199] | 0.30 ± 0.23 [0.195] |
| Elo-rating | -0.24 ± 0.11 [0.032] | -0.16 ± 0.11 [0.152] | -0.13 ± 0.12 [0.286] | -0.22 ± 0.12 [0.054] |
| Species (Stro)** | 0.65 ± 0.13 [<0.001] | **0.65 ± 0.13 [<0.001]** | **0.65 ± 0.13 [<0.001]** | **0.65 ± 0.13 [<0.001]** |
| Species (Tri)*** | -0.67 ± 0.12 [<0.001] | **-0.67 ± 0.12 [<0.001]** | **-0.67 ± 0.12 [<0.001]** | **-0.67 ± 0.12 [<0.001]** |
| Random effects | *Variance* | | | |
| Collection Date | 0.02 | 0.02 | 0.02 | 0.02 |
| Individual ID | 0.27 | 0.24 | 0.26 | 0.30 |

** Sex (male) shows model coefficients for males compared with the reference value, females*
*** Species (Stro) shows model coefficients for the parasite species S. fuelleborni compared with the reference value, O. aculeatum*
**** Species (Tri) shows model coefficients for the parasite species T. trichiura compared with the reference value, O. aculeatum*

Table SI-5. Results of GLMMs testing for variation in geohelminths infection intensity (EPG) among **adult female** Japanese macaques on Koshima island. LRT refers to likelihood ratio test, here shown with Chi-square $\chi^2$ tests, degrees of freedom (d.f.) and p-values (p). Estimates (β) are given with standard errors (SE) and p-values (p). Statistically significant ($p < 0.05$) results from models that outperform the null model are set in bold text.

| *Centrality metric of focus* | *Degree* | *Strength* | *Eigenvector* | *Null* |
|---|---|---|---|---|
| *LRT against null* ($\chi^2$; d.f.; p) | 3.616; 2; 0.164 | **7.424; 2; 0.024** | 4.214; 2; 0.122 | / |
| Fixed effects | Estimate β ± SE [p] | | | |
| Intercept | 6.76 ± 0.14 [<0.001] | 6.76 ± 0.13 [<0.001] | 6.76 ± 0.14 [<0.001] | 6.76 ± 0.16 [<0.001] |
| Centrality metric | 0.25 ± 0.13 [0.045] | **0.32 ± 0.12 [0.006]** | 0.27 ± 0.13 [0.036] | / |
| Age | -0.02 ± 0.14 [0.885] | -0.09 ± 0.13 [0.477] | -0.11 ± 0.15 [0.438] | -0.04 ± 0.16 [0.786] |
| Elo-rating | -0.15 ± 0.14 [0.279] | -0.06 ± 0.13 [0.622] | -0.04 ± 0.14 [0.777] | -0.10 ± 0.15 [0.510] |
| Species (Stro)* | -1.11 ± 0.28 [<0.001] | **-1.00 ± 0.29 [<0.001]** | -1.09 ± 0.28 [<0.001] | **-1.10 ± 0.29 [<0.001]** |
| Species (Tri)** | -1.11 ± 0.18 [<0.001] | **-1.16 ± 0.18 [<0.001]** | -1.13 ± 0.18 [<0.001] | **-1.12 ± 0.18 [<0.001]** |
| Random effects | *Variance* | | | |
| Collection Date | 0.00 | 0.00 | 0.00 | 0.00 |
| Individual ID | 0.20 | 0.15 | 0.19 | 0.27 |

** Species (Stro) shows model coefficients for the parasite species S. fuelleborni compared with the reference value, O. aculeatum*

*** Species (Tri) shows model coefficients for the parasite species T. trichiura compared with the reference value, O. aculeatum*

Table SI-6. Results of GLMMs testing for variation in geohelminths infection intensity (EPG) among **juvenile** Japanese macaques on Koshima island. LRT refers to likelihood ratio test, here shown with Chi-square $\chi^2$ tests, degrees of freedom (d.f.) and p-values (p). Estimates (β) are given with standard errors (SE) and p-values (p). Statistically significant ($p < 0.05$) results from models that outperform the null model are set in bold text.

| *Centrality metric of focus* | *Degree* | *Strength* | *Eigenvector* | *Null* |
|---|---|---|---|---|
| *LRT against null* ($\chi^2$; d.f.; p) | 0.120; 2; 0.942 | 2.491; 2; 0.288 | 1.643; 2; 0.440 | / |
| **Fixed effects** | *Estimate β* ± SE [p] | | | |
| Intercept | 7.04 ± 0.16 [<0.001] | 7.03 ± 0.15 [<0.001] | 7.03 ± 0.16 [<0.001] | 7.05 ± 0.16 [<0.001] |
| Centrality metric | 0.03 ± 0.10 [0.763] | 0.13 ± 0.09 [0.169] | 0.12 ± 0.11 [0.255] | / |
| Age | -0.36 ± 0.10 [<0.001] | -0.39 ± 0.09 [<0.001] | -0.40 ± 0.10 [<0.001] | -0.37 ± 0.10 [<0.001] |
| Sex (male) | 0.46 ± 0.20 [0.026] | 0.46 ± 0.19 [0.015] | 0.46 ± 0.19 [0.017] | 0.44 ± 0.20 [0.026] |
| Elo-rating | -0.14 ± 0.10 [0.177] | -0.10 ± 0.10 [0.286] | -0.07 ± 0.11 [0.532] | -0.13 ± 0.10 [0.189] |
| Species (Stro)** | 1.10 ± 0.13 [<0.001] | 1.11 ± 0.13 [<0.001] | 1.11 ± 0.13 [<0.001] | 1.10 ± 0.13 [<0.001] |
| Species (Tri)*** | -0.28 ± 0.14 [0.052] | -0.27 ± 0.14 [0.057] | -0.27 ± 0.14 [0.062] | -0.28 ± 0.14 [0.051] |
| **Random effects** | *Variance* | | | |
| Collection Date | 0.05 | 0.05 | 0.05 | 0.05 |
| Individual ID | 0.13 | 0.11 | 0.12 | 0.13 |

** Sex (male) shows model coefficients for males compared with the reference value, females*
*** Species (Stro) shows model coefficients for the parasite species S. fuelleborni compared with the reference value, O. aculeatum*
**** Species (Tri) shows model coefficients for the parasite species T. trichiura compared with the reference value, O. aculeatum*

Table SI-7. Results of GLMMs testing for variation in geohelminths infection intensity (EPG) among **adult female** Japanese macaques on Koshima island, keeping centrality metrics calculated from the whole-group network (**without recalculation**). LRT refers to likelihood ratio test, here shown with Chi-square $\chi^2$ tests, degrees of freedom (d.f.) and p-value (p). Estimates (β) are given with standard errors (SE) and p-value (p). Statistically significant ($p < 0.05$) results from models that outperform the null model are set in bold text.

| *Centrality metric of focus* | *Degree* | *Strength* | *Eigenvector* | *Null* |
|---|---|---|---|---|
| *LRT against null* ($\chi^2$; d.f.; p) | 3.970; 2; 0.137 | **7.385; 2; 0.025** | 5.746; 2; 0.057 | / |
| Fixed effects | Estimate β ± SE [p] | | | |
| Intercept | 6.77 ± 0.15 [<0.001] | 6.76 ± 0.14 [<0.001] | 6.76 ± 0.14 [<0.001] | 6.76 ± 0.16 [<0.001] |
| Centrality metric | 0.22 ± 0.13 [0.078] | **0.36 ± 0.17 [0.031]** | 0.15 ± 0.14 [0.278] | / |
| Age | -0.07 ± 0.15 [0.640] | -0.29 ± 0.18 [0.118] | -0.12 ± 0.17 [0.491] | -0.04 ± 0.16 [0.786] |
| Elo-rating | -0.12 ± 0.14 [0.412] | -0.09 ± 0.14 [0.496] | -0.05 ± 0.16 [0.755] | -0.10 ± 0.15 [0.510] |
| Species (Stro)* | -1.12 ± 0.28 [<0.001] | **-1.08 ± 0.29 [<0.001]** | -1.12 ± 0.29 [<0.001] | -1.10 ± 0.29 [<0.001] |
| Species (Tri)** | -1.10 ± 0.18 [<0.001] | **-1.13 ± 0.18 [<0.001]** | -1.13 ± 0.18 [<0.001] | -1.12 ± 0.18 [<0.001] |
| Random effects | *Variance* | | | |
| Collection Date | 0.00 | 0.00 | 0.00 | 0.00 |
| Individual ID | 0.21 | 0.20 | 0.25 | 0.27 |

** Species (Stro) shows model coefficients for the parasite species S. fuelleborni compared with the reference value, O. aculeatum*
*** Species (Tri) shows model coefficients for the parasite species T. trichiura compared with the reference value, O. aculeatum*

Table SI-8. Results of GLMMs testing for variation in geohelminths infection intensity (EPG) among **juvenile** Japanese macaques on Koshima island, keeping centrality metrics calculated from whole-group network (**without recalculation**). LRT refers to likelihood ratio test, here shown with Chi-square $\chi^2$ tests, degrees of freedom (d.f.) and p-value (p). Estimates (β) are given with standard errors (SE) and p-value (p). Statistically significant ($p < 0.05$) results from models that outperform the null model are set in bold text.

| *Centrality metric of focus* | *Degree* | *Strength* | *Eigenvector* | *Null* |
|---|---|---|---|---|
| *LRT against null* ($\chi^2$; d.f.; p) | 0.332; 2; 0.847 | 3.621; 2; 0.164 | 1.324; 2; 0.516 | / |
| Fixed effects | *Estimate β* ± SE [p] | | | |
| Intercept | 7.05 ± 0.16 [<0.001] | 7.05 ± 0.16 [<0.001] | 7.06 ± 0.15 [<0.001] | 7.05 ± 0.16 [<0.001] |
| Centrality metric | -0.01 ± 0.10 [0.890] | 0.03 ± 0.09 [0.713] | 0.11 ± 0.10 [0.264] | / |
| Age | -0.38 ± 0.10 [<0.001] | -0.37 ± 0.10 [<0.001] | -0.35 ± 0.10 [<0.001] | -0.37 ± 0.10 [<0.001] |
| Sex (male)* | 0.43 ± 0.20 [0.032] | 0.43 ± 0.20 [0.028] | 0.41 ± 0.19 [0.032] | 0.44 ± 0.20 [0.026] |
| Elo-rating | -0.13 ± 0.11 [0.234] | -0.13 ± 0.10 [0.201] | -0.09 ± 0.10 [0.362] | -0.13 ± 0.10 [0.189] |
| Species (Stro)** | 1.10 ± 0.13 [<0.001] | 1.10 ± 0.13 [<0.001] | 1.10 ± 0.13 [<0.001] | 1.10 ± 0.13 [<0.001] |
| Species (Tri)*** | -0.28 ± 0.14 [0.051] | -0.28 ± 0.14 [0.052] | -0.27 ± 0.14 [0.061] | -0.28 ± 0.14 [0.051] |
| Random effects | *Variance* | | | |
| Collection Date | 0.05 | 0.05 | 0.05 | 0.05 |
| Individual ID | 0.13 | 0.13 | 0.12 | 0.13 |

** Sex (male) shows model coefficients for males compared with the reference value, females*
*** Species (Stro) shows model coefficients for the parasite species S. fuelleborni compared with the reference value, O. aculeatum*
**** Species (Tri) shows model coefficients for the parasite species T. trichiura compared with the reference value, O. aculeatum*

Figure SI-9. **Results of GLMMs including centrality measures retained from networks where 5%, 10%, 25% or 50% of individuals were randomly removed.** Dark vertical lines represent the original result with dashed vertical lines representing its 95% confidence intervals. Dots and grey lines represent the knock-out results and their 95% confidence intervals. Red dots reflect statistically significant results (results that had a confidence interval not overlapping with 0).

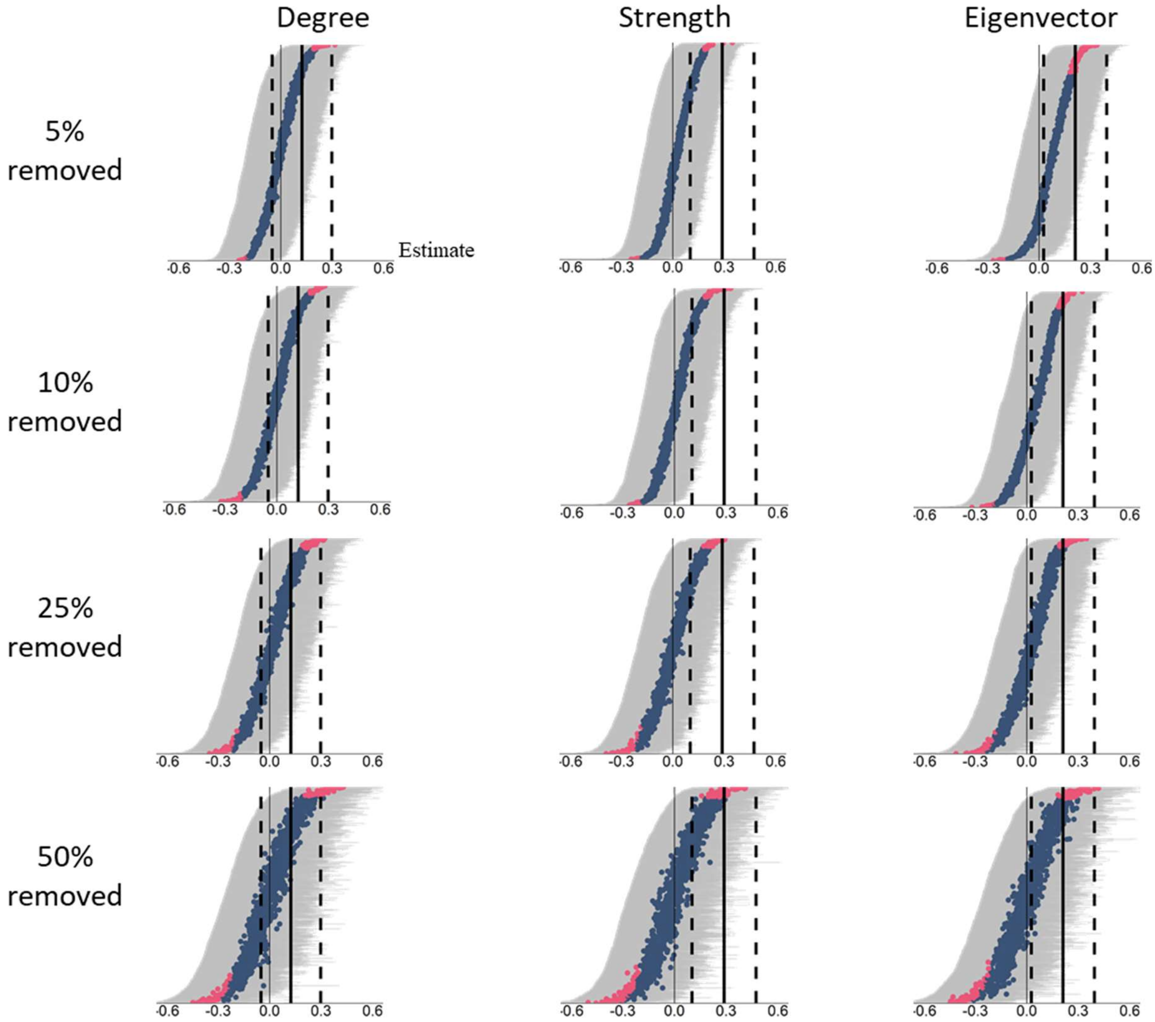

Table SI-10: **Results of 1000 random removals of individuals (% given in the first column) ran with centrality measures retained from the whole-group network (without recalculation)** and used to (re)model the relationship between network centrality and infection intensity. The heading "Number of model results equivalent to observed" indicates the number of simulation model results out of 1000 simulations that gave an equivalent result to models based on the whole-group network, i.e. for strength and eigenvecetor, positive estimates and 95% confidence intervals not overlapping zero, or for degree, a near-zero estimate with 95% confidence intervals overlapping zero.

| *Centrality measure* | Percentage of individuals removed | Number of model results equivalent to observed | Number of significant model results in the opposite direction |
|---|---|---|---|
| Degree | 5% | 96.3% | / |
| | 10% | 93.8% | / |
| | 25% | 87.8% | / |
| | 50% | 83.5% | / |
| Strength | 5% | 100% | 0% |
| | 10% | 99.1% | 0% |
| | 25% | 89.1% | 0% |
| | 50% | 55.5% | 0% |
| Eigenvector | 5% | 80.8% | 0% |
| | 10% | 68.7% | 0% |
| | 25% | 47.1% | 0% |
| | 50% | 35.0% | 0% |

Figure SI-11. **Results of GLMMs where 5%, 10%, 25% or 50% of individuals were randomly removed, but ran with centrality measures retained from the whole-group network (without recalculation)**. Dark vertical lines represent the original result with dashed vertical lines representing its 95% confidence intervals. Dots and grey lines represent the knock-out results and their 95% confidence intervals. Red dots reflect statistically significant results (results that had a confidence interval not overlapping with 0).

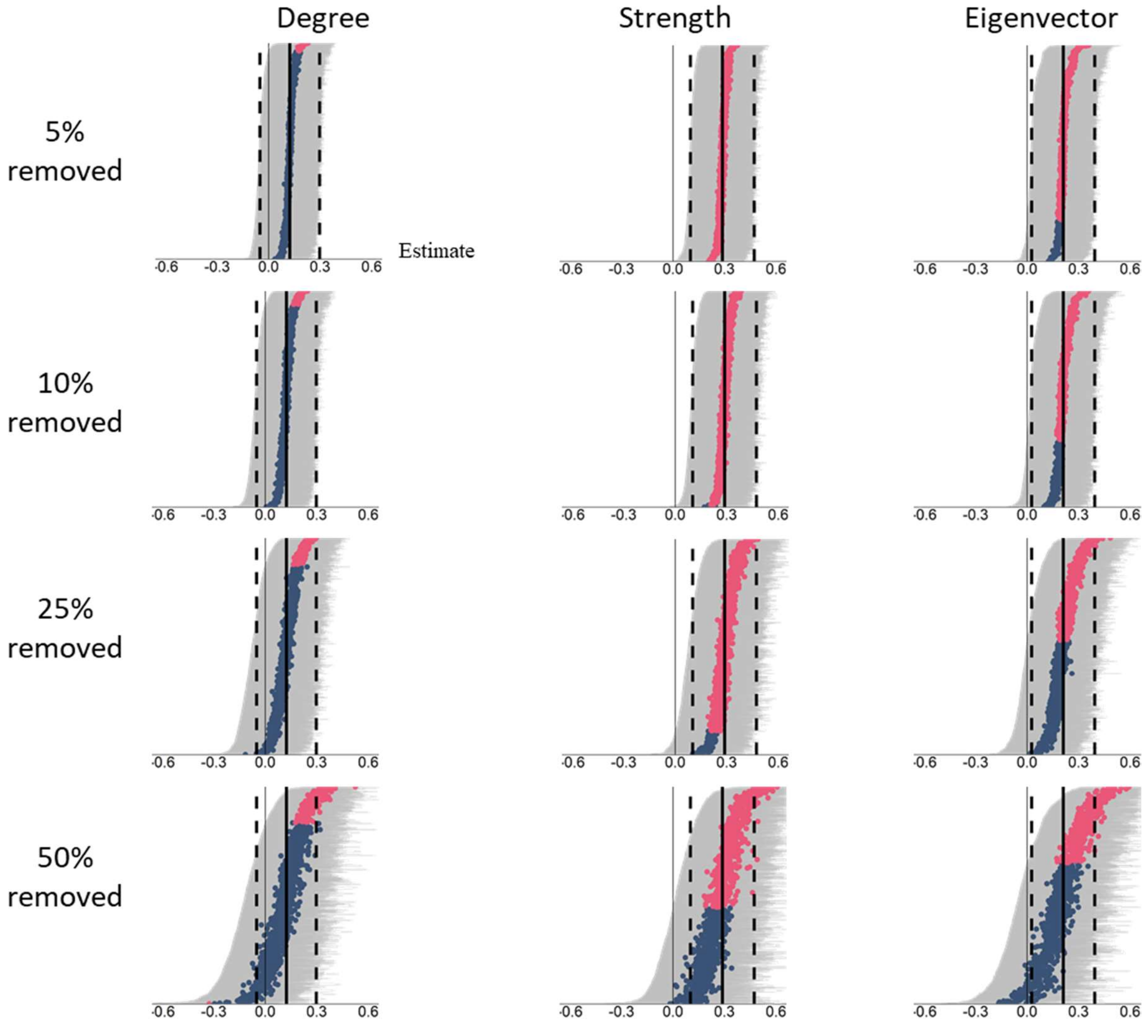
